# Immune and psychogenic fever arise through UCP1-independent thermogenic mechanisms

**DOI:** 10.64898/2026.01.30.702725

**Authors:** Takashi Matsuwaki, Wakana Harigai, Vladimir Maksimov, Kazuyuki Uchida, James K. Chambers, Keitaro Yamanouchi, Anna Eskilsson, Anders Blomqvist

## Abstract

Brown adipose tissue (BAT) thermogenesis is essential for cold defense, but its contribution to fever and emotionally induced hyperthermia (psychogenic fever) remains disputed. Here we address this issue using genetic, surgical, physiological, and molecular approaches in rats. We generated UCP1 knockout rats, in which classical BAT thermogenesis is abolished, and examined body temperature responses to systemic inflammation induced by lipopolysaccharide and to emotional stressors such as restraint and cage exchange. Despite profound impairment of cold-induced and β_3_-adrenergic-induced thermogenesis, UCP1 deletion did not affect LPS- or stress-evoked elevations in core or interscapular temperature. Surgical removal of interscapular BAT in wild-type rats likewise failed to alter these hyperthermic responses. Consistent with these findings, LPS and emotional stress induced only small, strain-dependent changes in the expression of thermogenic genes in BAT and minimally affected BAT mass, in marked contrast to the robust BAT activation elicited by β_3_-adrenergic stimulation. Notably, emotional stress induced UCP3 expression in neck muscles, suggesting a potential contribution of skeletal muscle metabolic processes to stress-induced hyperthermia. Together, these findings demonstrate that both immune-induced and psychogenic fever occur independently of BAT thermogenesis and point to non-BAT tissues - likely including skeletal muscle - as candidate peripheral effectors supporting fever and emotional hyperthermia.

## Introduction

While it is well established that brown adipose tissue (BAT) thermogenesis is critical for cold defense^1^, its role in fever and emotionally induced hyperthermia (psychogenic fever) remains disputed. Evidence from studies in rats has suggested that BAT thermogenesis may contribute to elevations in body temperature elicited by immune stimulation or emotional stress. For example, injection of prostaglandin E_2_ — the final mediator of inflammation-induced fever^2^ — into the thermoregulatory region of the preoptic hypothalamus not only raises body temperature but also increases sympathetic activity in sympathetic nerve bundles at the surface of interscapular BAT pads^3,4^. Such findings have been interpreted to suggest that the febrile response may be mediated through sympathetic activation of BAT. Additional support for this idea comes from observations that the increasein temperature within the BAT region precedes the rise in core body temperature following immune stimulation or during emotional stress^5–7^, consistent with — but not definitive for — a causal role for BAT thermogenesis.

However, these lines of evidence are indirect. In contrast, functional studies in mice lacking uncoupling protein 1 (UCP1) — the central mediator of BAT thermogenesis^1^ — showed no impairment of fever or emotional stress–induced hyperthermia^8–11^. Moreover, immune stimulation or emotional stress did not induce upregulation of UCP1 or associated thermogenic genes in wild-type animals, in stark contrast to the robust induction observed during cold exposure or following administration of a β_3_-adrenergic agonist^11^, both of which strongly activate BAT thermogenesis.

To resolve these conflicting findings — including the possibility of species differences — we generated UCP1 knockout rats and examined their temperature responses to immune stimulation induced by intraperitoneal injection of lipopolysaccharide, as well as to emotional stressors such as restraint or exposure to the scent of a conspecific male (cage exchange). Core and interscapular temperatures were recorded by telemetry using transponders implanted intraperitoneally or subcutaneously, respectively. In addition, interscapular BAT was surgically removed in wild-type rats. We find that neither UCP1 nor interscapular BAT is required for the elevation of body temperature during immune or emotional challenge, implying that the mechanisms governing thermoregulation during immune or emotional stress differ fundamentally from those that operate during cold exposure.

## Results

### Characterization of UCP1 knockout rats

Inactivation of the *Ucp1* gene, achieved by a targeted deletion in exon 2 (Fig. 1A), using the i-GONAD method^12^ and the CRISPR/Cas12 system, was verified by PCR and Western blot analyses, which confirmed the absence of *Ucp1* mRNA and UCP1 protein, respectively (Fig. 1B,C).

**Fig. 1.**
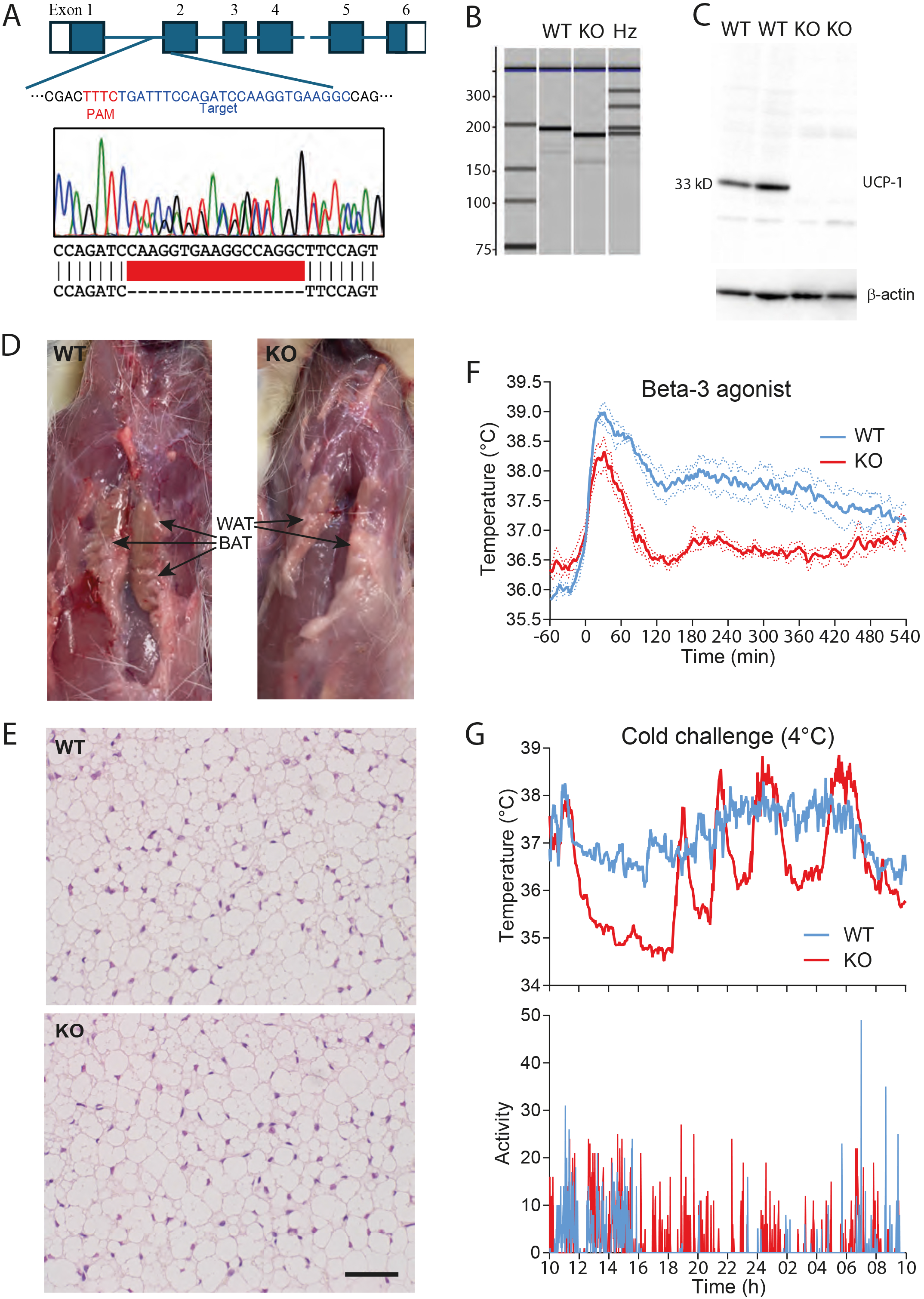
Generation and characterization of UCP1 knockout rats. *A*, Strategy for targeted deletion of the *Ucp1* gene. Top: The target sequence (blue) was TGATTTCCAGATCCAAGGTGAAGGC, adjacent to a TTTC PAM site (red) in exon 2. Bottom: Site of the deletion (red bar) as determined by sequencing. *B*, Microchip-based capillary electrophoresis showing a 17-bp deletion in the rat *Ucp1* gene; the two upshifted bands in heterozygous (Hz) rats represent heteroduplexes. *C*, Western blot confirming the absence of UCP1 protein in UCP1 knockout (KO) rats. *D*, Interscapular brown adipose tissue (BAT) present in wild-type (WT) rats is replaced by white adipose tissue (WAT) in UCP1 KO rats. *E*, Hematoxylin–eosin staining of interscapular adipose tissue revealing enlarged lipid vacuoles in UCP1 KO rats. Scale bar, 50 µm. *F*, A β_3_-adrenergic agonist induces sustained hyperthermia in WT but not UCP1 KO rats; the initial temperature peak reflects handling stress associated with injection at time 0. Solid lines show the mean, and dotted lines show the SEM. *n* = 4. *G*, UCP1 KO rats display impaired cold defense during exposure to 4 °C.

UCP1 knockout (KO) rats exhibited normal gross development. However, body weight in male KO rats was slightly lower than in wild-type (WT) or heterozygous (Hz) rats, which did not differ from each other (one-way ANOVA: F_2,24_ = 3.597, *P* = 0.0430; *P* < 0.05 for WT and Hz vs. KO). Female KO rats showed a tendency toward lower body weight between 5 and 9 weeks of age, but this difference was no longer present by 11 weeks (Suppl. Fig. 1).

Adipose tissue in the interscapular region — where BAT is located in WT rats — appeared whitish in KO rats (Fig. 1D). Histological analysis revealed enlarged lipid vacuoles (Fig. 1E), consistent with lipid accumulation resulting from the absence of UCP1-mediated uncoupling.

Administration of the β_3_-adrenergic agonist CL 316243 (0.5 mg/kg, i.p.), a potent inducer of BAT thermogenesis^13^, elicited a similar initial handling stress-induced hyperthermia in both WT and UCP1 KO rats. In WT rats, body temperature remained elevated following injection, whereas in KO rats it returned to pre-injection levels (two-way ANOVA for 120-540 min: *F*_1,6_ = 14.7, *P* = 0.0095), indicating that the β_3_-agonist failed to induce thermogenesis in the absence of UCP1 (Fig. 1F).

Consistent with impaired BAT function, UCP1 KO rats exhibited abnormal cold defense. When exposed to an ambient temperature of 4°C, WT rats maintained a stable body temperature with a normal diurnal rhythm, characterized by slightly lower temperatures during the light (inactive) period than during the dark (active) period. In contrast, UCP1 KO rats displayed a lower and more variable baseline body temperature. Periods of increased locomotor activity were observed, corresponding to transient elevations in body temperature (Fig. 1G; Suppl. Fig. 2).

### Emotional stress and immune challenge induce comparable increases in core and interscapular temperatures in UCP1 knockout and wild-type rats

As shown in Fig. 1F, UCP1 KO rats exhibited handling stress-induced hyperthermia comparable to that observed in WT rats following intraperitoneal injection. We next examined thermoregulatory responses to emotional stress using two other paradigms: cage exchange and restraint stress. In separate experiments, we measured either intraperitonal (core) temperature or temperature subcutaneously in the interscapular region overlying the BAT.

Exposure to a cage previously occupied by another rat elicited pronounced hyperthermia in both UCP1 KO and WT rats (Fig. 2A_1_). A similar increase was observed in the interscapular region in both genotypes (Fig. 2A_1_). Likewise, restraint stress induced by confinement in a plastic cylinder produced a rapid increase in both intraperitonal temperature and interscapular temperature in UCP1 KO and WT rats (Fig. 2A_2_). Intraperitoneal injection of lipopolysaccharide (LPS), a well-established model for peripheral inflammation and fever^14,15^, produced, after the initial similar handling stress induced hyperthermia, a fever response that did not differ between genotypes (Fig. 2A_3_). Furthermore, both WT and KO rats showed a similar temperature increase in the interscapular region (Fig. 2A_3_). These results indicate that emotional stress- and immune-induced elevations in core body temperature and interscapular temperature occur independently of UCP1-mediated BAT thermogenesis.

**Fig. 2.**
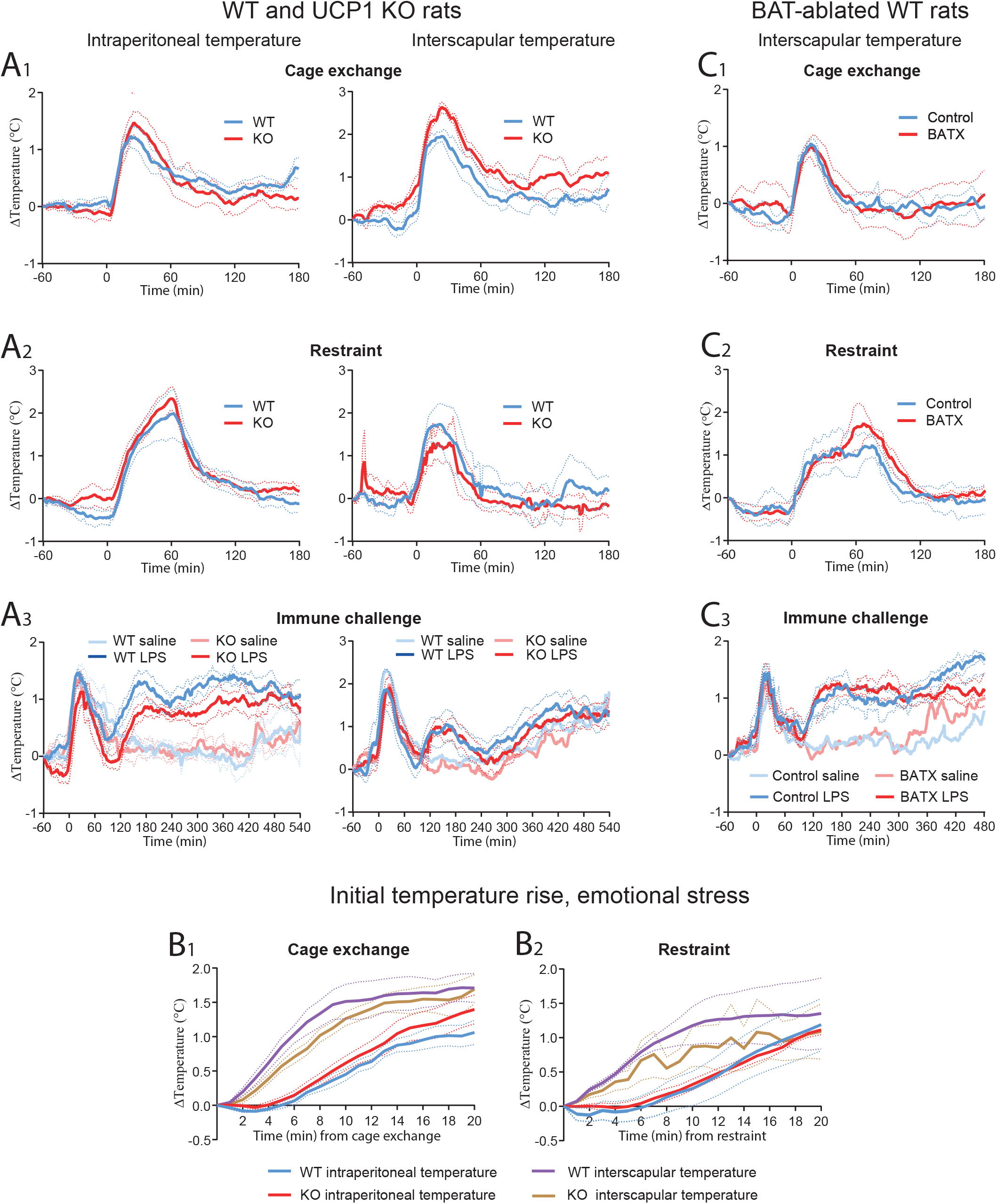
Temperature responses to emotional and immune stress. *A*, Intraperitoneal (left) and interscapular (right) temperature recordings in WT and UCP1 knockout (KO) rats during emotional stress. *B*, Interscapular temperature responses in WT rats with intact interscapular BAT (control) or following surgical removal of interscapular BAT (BATX). *(A1, B1)* Cage exchange; *(A2, B2)* Restraint; *(A3, B3)* Immune challenge with LPS. *C*, Initial temperature rise during (*C1*) cage exchange and (*C2*) restraint. All interventions were done at time point 0. Solid lines show mean, and dotted lines SEM. *n* = 3-5 for abdominal temperatures, and 3-4 for interscapular temperatures.

We next compared the temporal dynamics of temperature changes intraperitoneally and in the interscapular region following emotional stress. After both cage exchange and restraint, interscapular temperature increased within minutes, whereas the rise in intraperitoneal temperature was delayed by several minutes (Fig. 2B_1, 2_). The initial rate of temperature increase also tended to be steeper in the interscapular region than intraperitoneally (Fig. 2B_1, 2_), although this difference reached statistical significance only in wild-type rats subjected to cage exchange (linear regression: *F*_1,76_ = 19.75, *P* < 0.001).

### Removal of interscapular BAT does not affect emotional stress- or immune-induced increases in interscapular temperature

The findings above suggest that the rise in interscapular temperature following emotional or immune stress is not mediated by UCP1-dependent thermogenesis. To exclude the possibility that UCP1-independent thermogenic mechanisms within BAT contribute to this temperature increase^16^, we surgically removed the interscapular BAT and subsequently measured temperature in this region using a subcutaneously implanted transponder.

There was no difference in the interscapular temperature response to restraint stress, cage exchange, or immune challenge with LPS between sham-operated rats and rats lacking interscapular BAT. All three stimuli elicited comparable fever or stress-induced hyperthermic responses regardless of BAT removal (Fig. 2C_1-3_).

### Immune stimulation and emotional stress induce little or no expression of thermogenic genes in BAT and minimally affect BAT mass

Immune challenge of Wistar–Imamichi rats with LPS (120 μg kg^−1^, i.p.) did not induce transcription of *Ucp1* or *Ppargc1a*—the latter encoding the transcriptional coactivator PGC-1α, a key regulator of genes involved in oxidative metabolism (Cannon & Nedergaard, 2004)— relative to saline-treated controls (Fig. 3A_1,2_). LPS also failed to significantly reduce BAT weight (Fig. 3A_3_) or alter BAT appearance (Fig. 3B). In contrast, administration of the β_3_-adrenergic agonist CL 316243 (1.0 mg kg^−1^, i.p.) robustly induced *Ucp1* and *Ppargc1a* transcription (one-way ANOVA: *Ucp1 F*_2,15_ = 14.66, *P* = 0.0003; *P* = 0.001 vs. saline; *Ppargc1a F*_2,15_ = 23.71, *P* < 0.001;*P* < 0.001 vs. saline) (Fig. 3A_1,2_), significantly reduced BAT weight (*F*_2,17_ = 6.334, *P* = 0.0088; *P* = 0.0066 vs. saline) (Fig. 3A_3_), and resulted in marked darkening of the BAT (Fig. 3B).

**Fig. 3.**
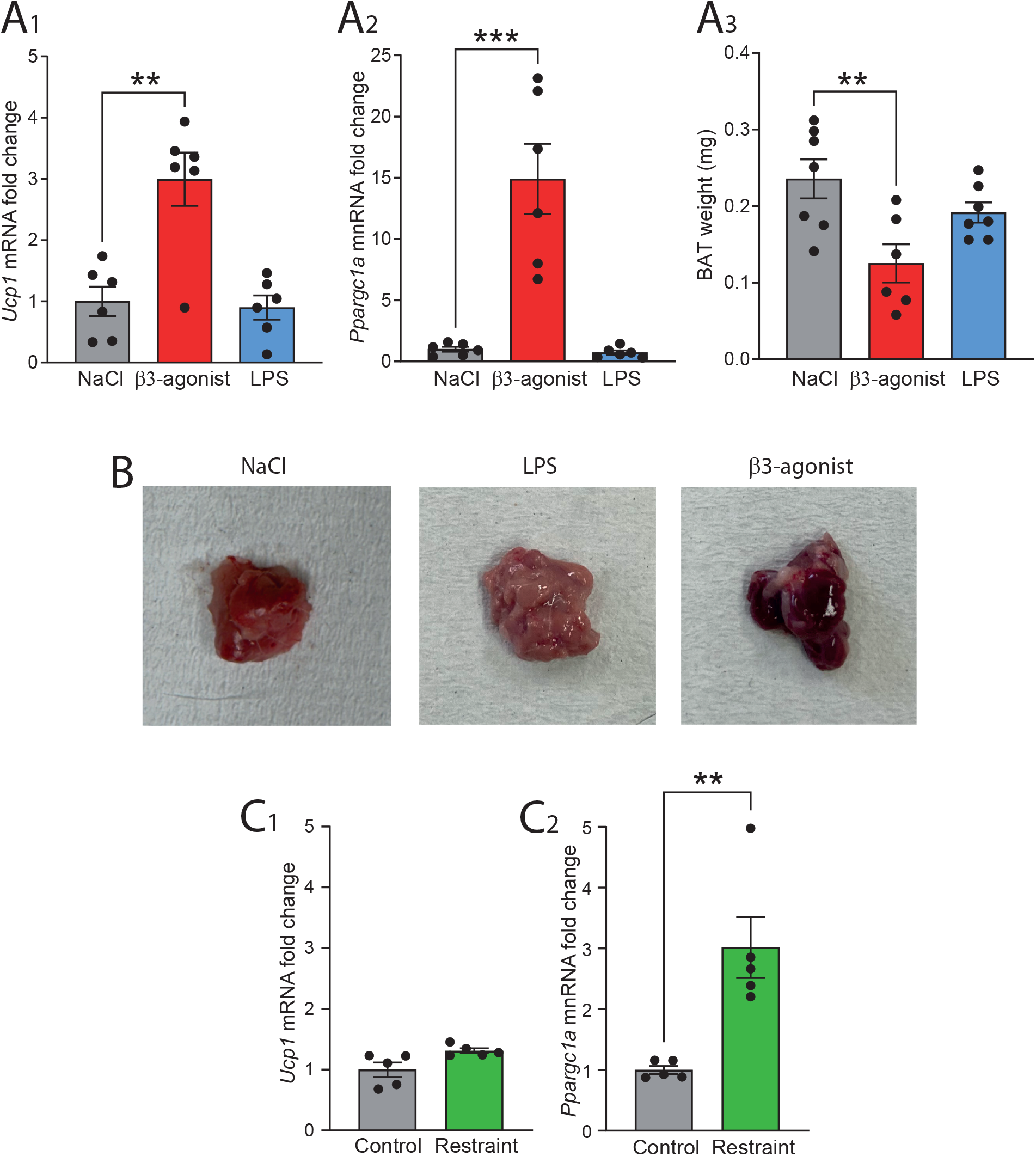
Effects of β_3_-adrenergic stimulation, immune challenge, and emotional stress on thermogenic gene expression and BAT morphology in WT rats. *A*. β_3_-adrenergic stimulation and immune challenge. *A1, Ucp1* mRNA expression. *A2, Ppargc1a* mRNA expression. *A3*, BAT weight. *n* = 6-7. *B*. Darkening of BAT following intraperitoneal β_3_-adrenergic agonist administration, but not after LPS. *C*. Restraint stress. *C1, Ucp1* mRNA expression. C2, *Ppargc1a* mRNA expression. Error bars indicate SEM. \**P* < 0.05; \*\**P* < 0.01;\*\*\**P* < 0.001.

In a separate experiment, we examined the effects of restraint stress on thermogenic gene expression in BAT, using naïve rats as controls. As shown in Fig. 3C, restraint produced only a minor, statistically non-significant increase in *Ucp1* expression. *Ppargc1a*, however, was significantly upregulated, although still at a low magnitude relative to β_3_-adrenergic stimulation.

In Sprague Dawley rats, LPS elicited small, statistically non-significant increases in *Ucp1* and *Ppargc1a* expression. In contrast, β_3_-agonist administration produced a robust induction of these genes (one-way ANOVA: *Ucp1 F*_3,20_ = 18.70, *P* < 0.001; *P* < 0.001 vs. saline; *Ppargc1a F*_3,20_ = 246.6, *P* < 0.001; *P* < 0.001 vs. saline), similar to the response observed in Wistar–Imamichi rats. Emotional stress induced by cage exchange likewise produced only modest increases in *Ucp1* and *Ppargc1a* expression (Suppl. Fig. 3A, B).

β_3_-agonist treatment significantly reduced BAT weight in Sprague Dawley rats (one-way ANOVA: *F*_3,20_ = 4.554, *P* = 0.0137; *P* = 0.0315 vs. saline). By contrast, LPS induced only a small, non-significant decrease in BAT weight, and cage exchange produced no detectable change (Suppl. Fig. 3C).

### Emotional stress induces *Ucp3* expression in neck muscles

Restraint stress significantly increased *Ucp3*, but not *Ucp2*, mRNA expression in neck muscles, consistent with our previous findings in mice and with UCP3 being a regulator of muscle energy metabolism (Della Guardia et al., 2024). One hour after release from restraint, *Ucp3* mRNA levels were elevated 1.6-fold relative to controls (*t*-test; *n* = 9–10; *P* = 0.0265), whereas *Ucp2* expression did not differ between groups (Fig. 4).

**Fig. 4.**
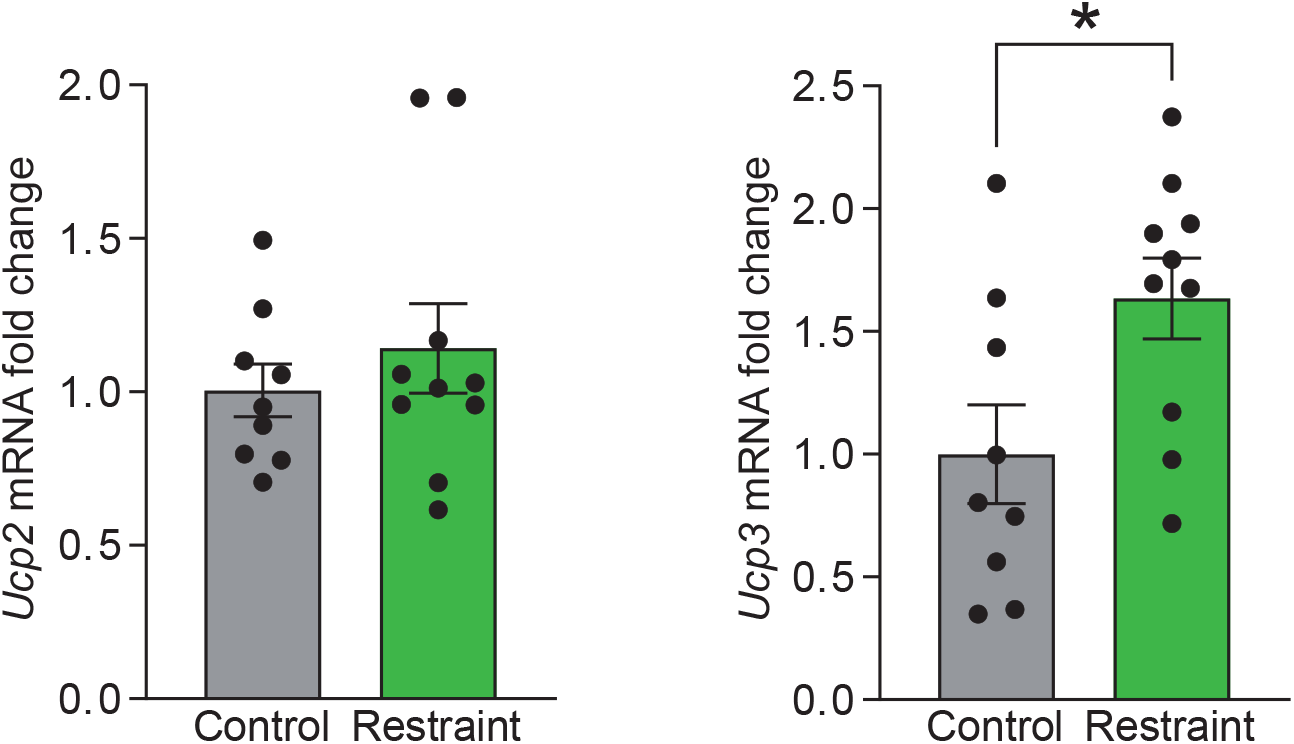
Effects of restraint stress on *Ucp2* and *Ucp3* mRNA expression in neck muscles. *A, Ucp2* mRNA expression. B, *Ucp3* mRNA expression. Error bars indicate SEM. \**P* < 0.05.

## Discussion

We here demonstrate in rats that fever induced by peripheral injection of lipopolysaccharide (LPS), a widely used model of systemic inflammation and fever^15^, as well as hyperthermia induced by emotional stress (psychogenic fever)^17^, occurs in the absence of UCP1, consistent with our previous observations in mice^11^. To the best of our knowledge, this study represents the first use of a UCP1 knockout rat, generated de novo for the present work, thereby establishing a novel genetic model to directly test the role of brown adipose tissue (BAT) thermogenesis in fever. Because UCP1 is indispensable for BAT thermogenesis^1,18^, the present findings provide direct genetic evidence that neither inflammation-induced fever nor psychogenic fever depends on classical BAT-mediated heat production. Collectively, our results challenge the prevailing view that central fever-generating circuits recruit the same peripheral thermogenic mechanisms as those engaged during cold exposure, namely uncoupled mitochondrial respiration in brown adipocytes^19,20^.

As outlined in the Introduction, the evidence supporting a critical role for BAT thermogenesis in immune-induced and psychogenic fever has been largely indirect. This includes reports of increased sympathetic nerve activity in fibers presumed to innervate BAT^3^, as well as observations that temperature in the interscapular region rises before core body temperature during immune challenge or emotional stress^5–7^. We here show, in consistency with previous observations, that the increase in interscapular temperature during emotional stress indeed occurs prior to the rise in intraperitoneal(core body) temperature, but most importantly, that the same phenomenon is seen also in *Ucp1* KO animals. The persistence of the interscapular temperature response following surgical removal of interscapular BAT further demonstrates that the temperature increase in the interscapular region cannot be attributed to BAT thermogenesis.

The present findings imply that increases in interscapular temperature during fever and emotional stress reflect mechanisms other than local heat production by BAT. Likely contributors include activation of neck and shoulder musculature and increased regional blood flow, both of which are known to accompany emotional stress^21^. It has been demonstrated that muscle temperature increases rapidly when rodents are exposed to the odor of a predator^22^, which is consistent with our observation that emotional stress induces *Ucp3* expression in neck muscles. UCP3 is highly expressed in skeletal muscle^23^, and while it does not mediate adaptive thermogenesis ^24^, it has been suggested to participate in metabolic or redox-regulatory mechanisms that support the energetic demands of hyperthermia^25,26^. Notably, also immune challenge results in increased *Ucp3* transcription in muscle^11,25^, and *Ucp3* deletion in skeletal muscle has been reported to abolish the febrile response^10^. The available data hence point to a model in which skeletal muscle metabolism, and not BAT thermogenesis, elicits immune- and emotional stress-induced increases in body temperature.

While the present study thus demonstrates that immune-induced and psychogenic fever can occur entirely independently of UCP1-mediated thermogenesis, our findings may nevertheless allow reconciliation with earlier reports suggesting BAT involvement. In contrast to our previous mouse studies, in which neither immune stimulation nor emotional stress induced *Ucp1* or *Ppargc1a* transcription^11^ (but cf. ref. ^27^), Sprague Dawley rats — but not Wistar–Imamichi rats — displayed a modest induction of *Ucp1* mRNA following LPS administration, and both rat strains showed modest *Ppargc1a* mRNA induction after emotional stress. In both rat strains, these transcriptional changes were accompanied by small, statistically non-significant reductions in BAT mass after immune challenge, whereas BAT mass remained unchanged after cage exchange in the strain examined (Sprague Dawley). Together, these observations suggest that limited UCP1-dependent BAT activation may occur during immune-induced fever (and possibly also after emotional stress), although it is clearly not required for the febrile response itself. It remains possible, however, that such auxiliary BAT activation becomes relevant during more prolonged or energetically demanding inflammatory states, when alternative energy reserves become constrained, as has recently been proposed (Li et al., 2024).

In summary, our findings demonstrate that fever and emotional stress–induced hyperthermia are fundamentally distinct from adaptive brown adipose tissue–mediated thermogenesis, at least under the experimental conditions examined here. Using complementary genetic, surgical, physiological, and molecular approaches, we show that neither UCP1 nor interscapular BAT is required for the elevation of body temperature during immune challenge or emotional stress. Instead, increases in interscapular temperature during these states likely reflect altered heat distribution and non-BAT metabolic processes, potentially involving skeletal muscle. Together, these results redefine fever as a UCP1-independent hyperthermic state driven by non-BAT thermogenic pathways, highlighting a clear mechanistic separation between thermoregulation during cold exposure and during immune or emotional stress.

## Methods

### Animals

Adult Wistar–Imamichi rats (Imamichi Institute for Animal Reproduction, Tsuchiura, Japan) and Sprague–Dawley rats (Janvier Labs, Le Genest-Saint-Isle, France) were used in this study.Animals were housed under controlled environmental conditions (22–23 °C) with a 14 h light/10 h dark cycle or a 12 h light/12 h dark cycle, respectively, and had ad libitum access to food and water.

All experimental procedures were approved by the Institutional Animal Care and Use Committee of the University of Tokyo and by the Animal Ethics Committee in Linköping and were conducted in accordance with international guidelines for the care and use of laboratory animals.

### Generation of UCP1 knockout rats

UCP1 knockout (KO) rats were generated using the improved Genome-editing via Oviductal Nucleic Acids Delivery (i-GONAD) method^12^. Genome editing was performed with the CRISPR/Cas12 system. The gRNA and Cas12 protein (Alt-R CRISPR-Cpf1 crRNA; Alt-R® A.s. CRISPR-Cas12a (Cpf1) V3) were obtained from Integrated DNA Technologies (Coralville, IA, USA). The target sequence was 5′-TGATTTCCAGATCCAAGGTGAAGGC-3′, adjacent to a TTTC PAM site in exon 2.

Female rats were anesthetized with isoflurane (Viatris, Tokyo, Japan) on the day after mating. The prepared genome-editing solution was injected into the oviductal ampulla using a glass pipette (Narishige, Tokyo, Japan). Electroporation was then performed three times using a CUY21EDIT device (Nepagene, Tokyo, Japan) at 60 V, with a pulse-on duration of 5.5 ms and a pulse-off interval of 50.0 ms.

Founder animals were backcrossed to the Wistar–Imamichi strain for two generations before experimental use. Wild-type littermates served as controls.

### Genotyping

Genotyping of offspring was performed by PCR using genomic DNA extracted from tail tissue. PCR reactions were carried out with KOD DNA polymerase (Toyobo, Osaka, Japan). A primer pair, UCP1-F (5′-TTATTTAGTCCCCTACCCACCC-3′) and UCP1-R (5′-CTCTTGGACCGTATCGTAGAGG-3′), was used to distinguish WT (263 bp) and KO (246 bp) alleles. The PCR protocol consisted of an initial denaturation at 94 °C for 2 min, followed by 35 cycles at 98 °C for 20 s, 60 °C for 30 s, and 68 °C for 30 s.

PCR products were visualized using a Microchip Electrophoresis System (MultiNA, Shimadzu Corporation, Kyoto, Japan) with the DNA-500 kit (Shimadzu).

### Western blot analysis

Proteins isolated from BAT were separated on 8% Tris–glycine gels, transferred to PVDF membranes, and immunoblotted with a rabbit anti–UCP1 antibody (1:2500; U6382, MilliporeSigma, Burlington, MA). Signals were detected using a horseradish peroxidase– conjugated anti-rabbit IgG antibody and ECL Prime (Amersham Biosciences, Buckinghamshire, UK). β-actin was used as a loading control and detected with a mouse anti-β-actin antibody (1:4000; Sigma, St. Louis, MO, USA).

### Histology

Interscapular adipose tissue was dissected, fixed in 4% paraformaldehyde, embedded in paraffin, and sectioned at 7 μm. Sections were stained with hematoxylin and eosin and imaged by light microscopy to assess tissue morphology and lipid content.

### Surgery

Animals were anesthetized with isoflurane and implanted with a temperature- and activity-sensing transponder (E-Mitter Telemetry System; STARR Life Sciences, Oakmont, PA).Transponders were placed either intraperitoneally to measure core body temperature or subcutaneously in the interscapular region to measure local temperature. After implantation, animals were housed individually.

In a subset of experiments, interscapular brown adipose tissue (BAT) was surgically removed prior to transponder implantation. All surgical procedures were performed under aseptic conditions, and animals were allowed to recover for 1 week before experimental testing.

### Cold stress, emotional stress, immune challenge, and adrenergic activation of BAT

#### Cold stress

One week after transponder implantation, animals were exposed to 4 °C for 1 week, during which body temperature was continuously monitored by telemetry.

#### Emotional stress, restraint stress, and immune challenge

Emotional stress was induced by transferring rats from their home cage to the cage of another unfamiliar rat (“cage exchange”). Two days later, restraint stress was applied by immobilizing the animals for 1 h in a plastic bag (DecapiCone, Braintree Scientific Inc., Braintree, MA). Four days after restraint stress, rats received an intraperitoneal injection of lipopolysaccharide (LPS; O111:B4; Sigma-Aldrich, St. Louis, MO; 120 μg kg−^1^) or saline. After a 5-day washout period, treatments were crossed over such that animals previously given LPS received saline and vice versa.

#### Adrenergic activation of BAT

Approximately 1 week after transponder implantation, rats were administered a β_3_-adrenergic agonist (SR59230A; Tocris Bioscience, Bristol, UK; 1.0 mg kg^−1^, i.p.) or saline.

All injections and stress-inducing procedures were performed at approximately 10:00 a.m. to minimize circadian variability.

### Quantitative real-time PCR

Rats were injected intraperitoneally with lipopolysaccharide (LPS; 120 µg kg^−1^), the β_3_-adrenergic agonist CL 316243 (1.0 mg kg^−1^), or saline, or were subjected to cage exchange or restraint stress as described above. They were euthanized by CO_2_ asphyxiation at defined time points: 4 h after LPS injection, 2 h after β_3_-agonist injection, 2–3 h after cage exchange, or 1 h after termination of restraint stress.

Interscapular BAT and neck muscles were rapidly excised, placed in RNAlater stabilization reagent (Qiagen, Hilden, Germany), and stored at −70 °C until further processing. Total RNA was extracted using the RNeasy Universal Plus kit (Qiagen), and cDNA was synthesized using the High-Capacity cDNA Reverse Transcription Kit (Applied Biosystems, Foster City, CA).

Quantitative real-time PCR was performed using TaqMan Gene Expression Master Mix (Applied Biosystems) on a 96-well plate with a QuantStudio™ 7 Flex Real-Time PCR System (Applied Biosystems). The following TaqMan Gene Expression Assays (Thermo Fisher Scientific, Stockholm, Sweden) were used: *Ucp1* (Rn00562126_m1), *Ucp2* (Rn01754856_m1), *Ucp3* (Rn00565874_m1), *Ppargc1a* (PGC-1α; Rn00580241_m1), and *Gapdh* (Rn01775763_g1).

Gene expression levels were calculated using the comparative Ct (ΔΔCt) method. Target gene expression was normalized to the *Gapdh* and expressed relative to the saline-treated control group.

### Data processing and statistics

Normalization of the temperature recordings was done by subtracting, for each animal, the temperature recorded at 60 min prior to injection/emotional stress, or, for analyses of initial response to emotional stress, the value recorded at time point 0.

Statistical analyses were performed using GraphPad Prism version 10 (GraphPad Software, San Diego, CA). Data are presented as mean ± SEM. Group differences were assessed using unpaired *t*-tests or, when three or more groups were compared, one-way or two-way ANOVA with correction for multiple comparisons by Tukey’s post hoc test or the two-stage linear step-up procedure of Benjamini, Krieger and Yekutieli, respectively. A *P* value < 0.05 was considered statistically significant.

## Author contributions

A.B., A.E., and T.M. conceived and designed the experiments. A.E., J.K.C., K.Y., U.H., and V.M. performed the experiments. A.B. and T.M. wrote the manuscript.

## Acknowledgement

This study was supported by JSPS KAKENHI Grant Nos. 22H00396 and 23KK0126 to T.M., and by the Swedish Brain Foundation (Grant FO2025-0110-HK-175) to A.B.

