## Supplementary figures and images for "Immune and psychogenic fever arise through UCP1-independent thermogenic mechanisms"

### Suppl. Figs 1-3

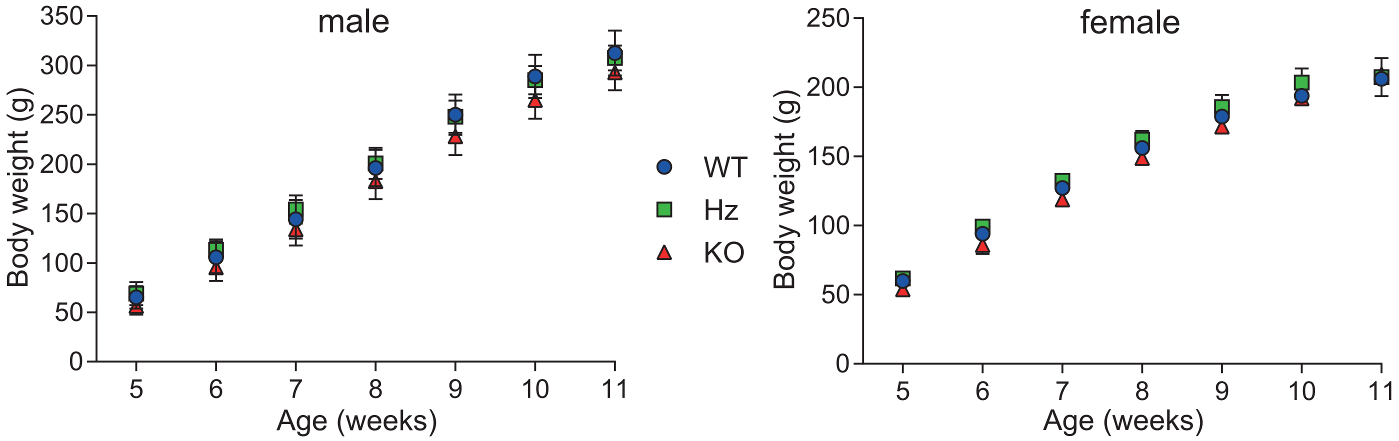

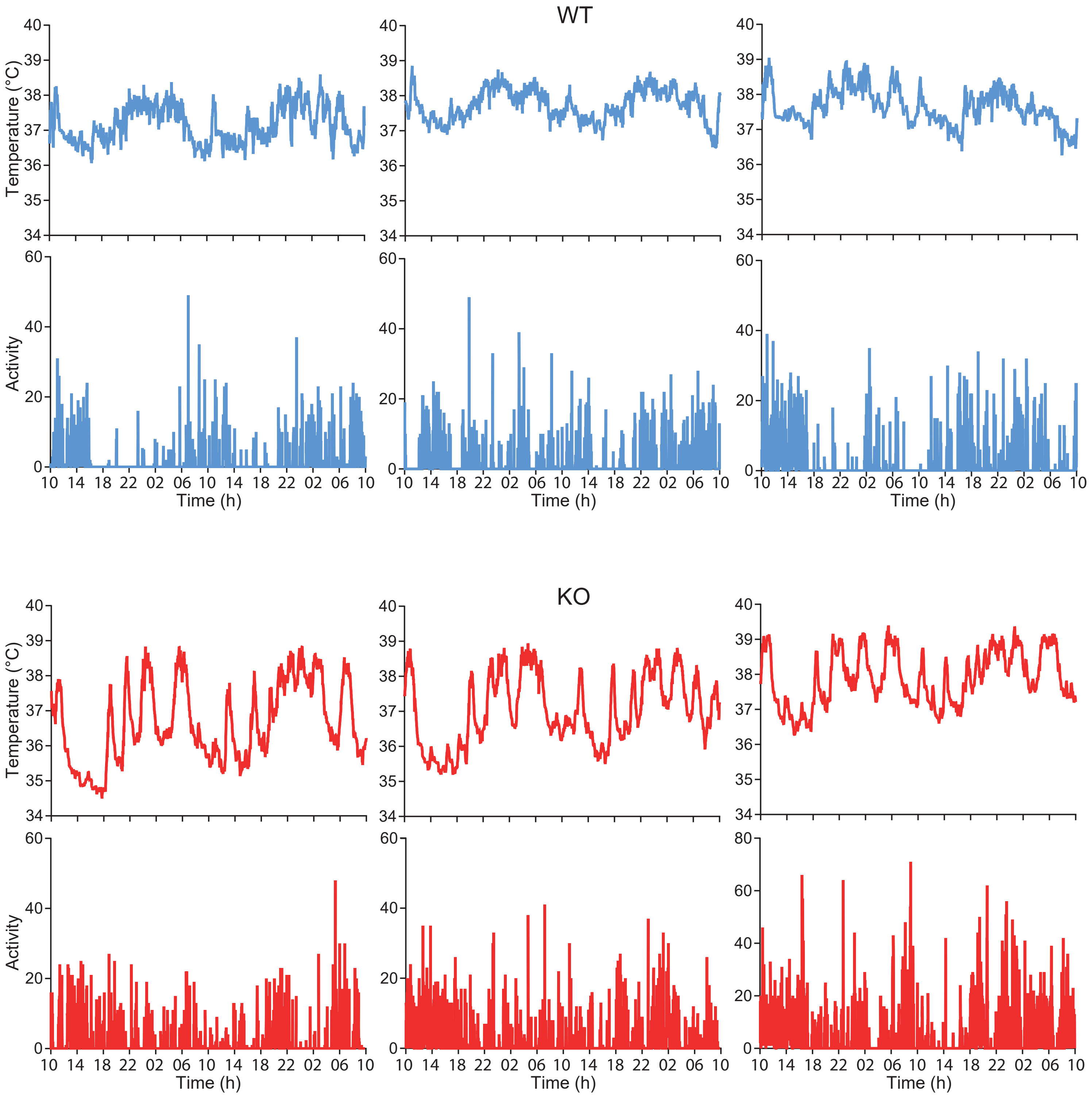

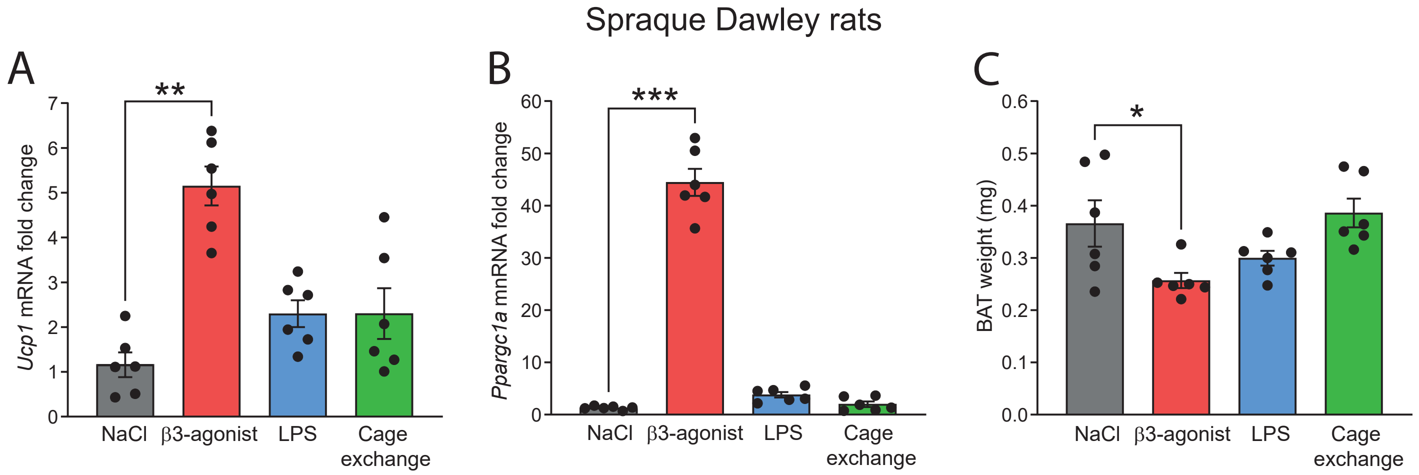
